# Probing nanomotion of single bacteria with graphene drums

**DOI:** 10.1101/2021.09.21.461186

**Authors:** I.E. Rosłoń, A. Japaridze, P.G. Steeneken, C. Dekker, F. Alijani

**Author notes:** Contributed equally to this work as first authors (I.E. Rosłoń, A. Japaridze). Corresponding author: F. Alijani.

## Abstract

Motion is a key characteristic of every form of life^1^. Even at the microscale, it has been reported that colonies of bacteria can generate nanomotion on mechanical cantilevers^2^, but the origin of these nanoscale vibrations has remained unresolved^3,4^. Here, we present a novel technique using drums made of ultrathin bilayer graphene, where the nanomotion of single bacteria can be measured in its aqueous growth environment. A single *E. coli* cell is found to generate random oscillations with amplitudes of up to 60 nm, exerting forces of up to 6 nN to its environment. Using mutant strains, we are able to pinpoint the bacterial flagella as the main source of nanomotion. By real-time tracing of changes in nanomotion upon administering antibiotics, we demonstrate that graphene drums can perform antibiotic susceptibility testing with single-cell sensitivity. These findings deepen our understanding of processes underlying cellular dynamics, and pave the way towards high throughput and parallelized rapid screening of the effectiveness of antibiotics in bacterial infections with graphene devices.

---

Living cells exhibit nanomechanical vibrations as a result of the biological processes that govern their growth, function, and reproduction^5^. This nanomotion is an intriguing phenomenon of unravelled origin that has been observed in a wide variety of living organisms, including neuronal cells^6^, erythrocytes, yeasts^7,8^, and bacteria^4^. Numerous hypotheses have been proposed for the underlying driving mechanism, such as motion of organelles, internal redistribution of cell membranes^9^ and the action of ion pumps^3^, but consensus has not been reached^4^. This relates to the fact that non-invasive probing of biomechanics at the microscale is highly challenging, which has stimulated the development and application of techniques like atomic force microscopy^10-12^ (AFM), optical and magnetic tweezers^13^, flow cytometry^14^, and optical tracking of cells^15,16^. In particular for bacterial cells, micromechanical cantilevers have emerged as powerful tools for detecting vibrations of adhered cell populations (100-1000 bacteria) in a liquid environment^4^. It was shown that the nanomotion of these populations rapidly decreases in the presence of antibiotics, which holds great promise for the development of rapid antibiotic susceptibility testing technologies^2^. Both for probing fundamental biomechanical processes and for development of nanomotion-based antibiotic susceptibility tests in medical diagnostics, it is crucial to elucidate the microscopic origins of nanomotion.

Here, we present a novel single-cell technique based on suspended graphene drums^17^, which greatly enhances the sensitivity of nanomechanical sensing compared to previous cantilever-based methods. The ultra-high sensitivity of the technique allowed us to clarify the mechanism that lies at the root of bacterial nanomotion by probing various strains of *Escherichia coli* (*E. coli*). The small mass, high stiffness, and micron-sized area of a suspended graphene drum enables detecting nanomotion at even the single bacterium level. Using arrays of these drums, we compare the vibrations produced by different *E. coli* strains. In particular, we investigate the contributions of the bacterial cell wall synthesis, flagella, rotor, and ion pump to nanomotion, and demonstrate that flagellar motion is the main source of nanomotion in these bacteria. Moreover, by tracing the nanomotion in the presence of antibiotics, we show that this novel ultrasensitive graphene-based platform enables antibiotic susceptibility tests with single-bacterium sensitivity. This opens new routes towards faster, label-free detection of antimicrobial resistance at the single-cell level with potential applications in drug screening and rapid diagnostics.

## Graphene drums for probing a single bacterium

The experiments were performed using drums made of ultrathin (<1 nm) bilayer CVD graphene that covered circular cavities with a diameter of 8 µm and a depth of 285 nm that were etched in SiO_2_. A silicon chip with an array of thousands of these graphene-covered cavities was placed inside a cuvette containing *E. coli* in Lysogeny Broth (LB) medium, where APTES was used to bind the bacteria to the graphene surface (see Supplementary Note 1 and Methods). The nanomotion of a bacterium resulted in a deflection of the suspended membrane, which was measured using laser interferometry^18^, see Figure 1a. The bacterium induced a time-dependent deflection *z*(*t*) at the center of the suspended graphene drum, which can be determined from the modulation of the intensity of the reflected light^19^. To quantitatively compare the nanomotion of different drums, we acquired *z*(*t*) traces over 30 second periods to obtain the variance *σ*^2^ = <*z*^2^(*t*)>, or the motion amplitude σ, which we used as a measure of the magnitude of the nanomotion.

**Figure 1.**
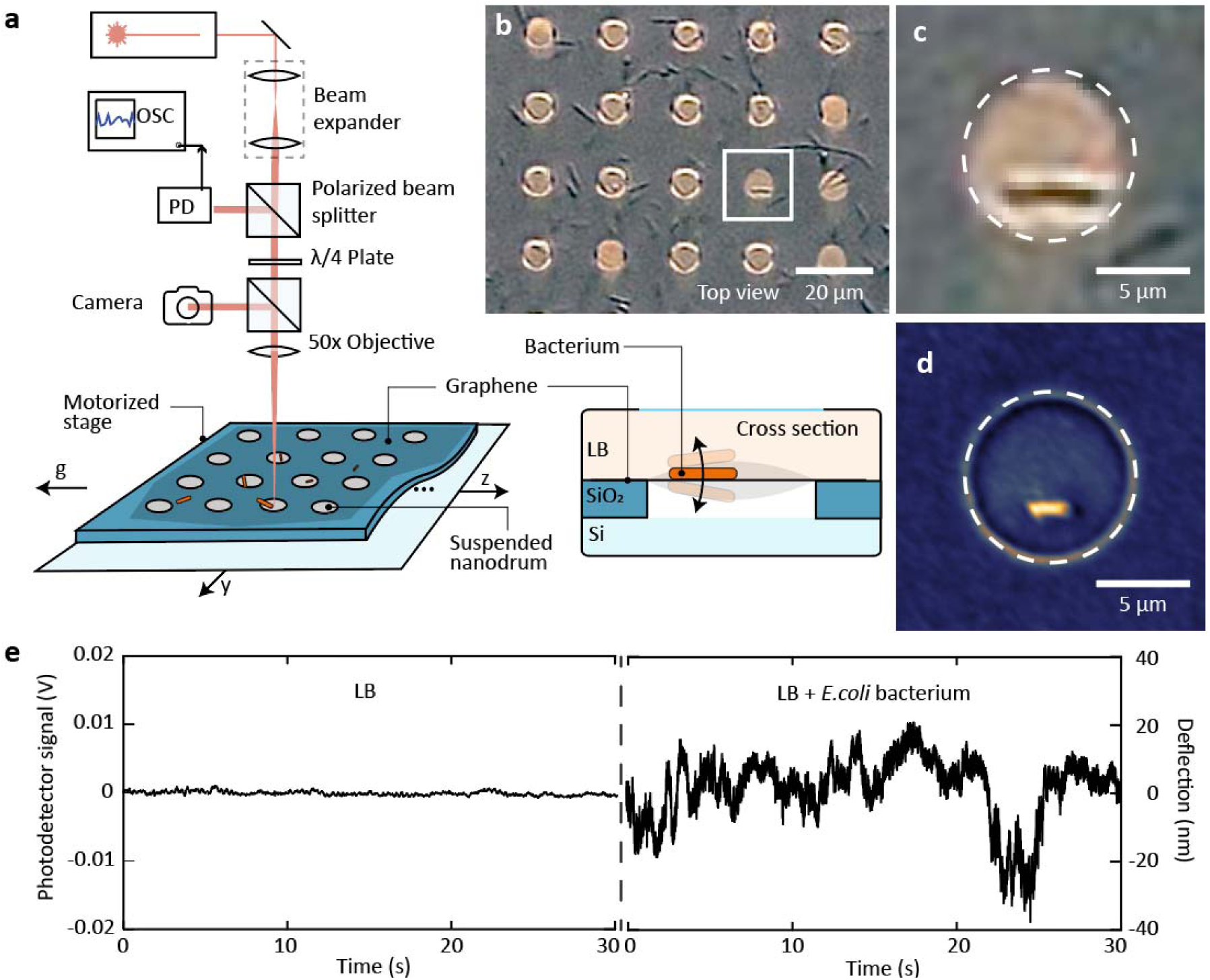
Detection of nanomotion of single bacteria by graphene drums. a) Schematic of the interferometric measurement setup used to record the nanomotion. b) Optical microscope image of an array of suspended drums with adhered *E. coli*. c) Zoom of the area indicated by a white square in panel b, showing a dividing bacterium on top of a graphene drum. d) SEM image of an *E. coli* on a suspended graphene drum. e) Recorded deflection of a suspended graphene drum immersed in LB without a bacterium (left), compared to the signal from a graphene drum with a bacterium present (right).

Drums containing a single live bacterium (Figure 1b-e) displayed large displacements *z*_max_ of up to 60 nm, with a time averaged motion amplitude of up to *σ* = 20 nm, that clearly exceed the deflection of drums without bacteria and signal from cells deposited on the Si/SiO_2_ substrate away from the drums, which yielded a background *σ* = 2 nm (see Figure 1f, Supplementary Notes 2 and 3). The large oscillation amplitudes can be associated with the movement of the suspended drum and originate from bacterial biophysical processes. To characterize the motion further, we recorded the signal of a single bacterium for more than 1 hour. It is apparent that fractal-like fluctuations were present over a wide range of timescales, see Figure 2a, where both low- and high frequency fluctuations are observed on timescales ranging from seconds to hours. Figure 2b displays the power spectral density of the motion (black line), compared to the background signal of an empty drum. The spectra show a 1/f^*α*^ frequency dependence with a mean value of *α* = 1.8 ± 0.1 (n=277 graphene drums; Figure 2c). The difference between drums with and without a single bacterium can also be clearly perceived by listening to audio recordings that were generated by converting the interferometric traces to a sound track (provided as Supplementary Audio). These results are consistent with power spectral densities found for bacterial colonies on AFM cantilevers^20^, and show that the nanomotion generated by even a single *E. coli* bacterium lacks a specific periodicity but instead involves a wide range of frequencies.

**Figure 2.**
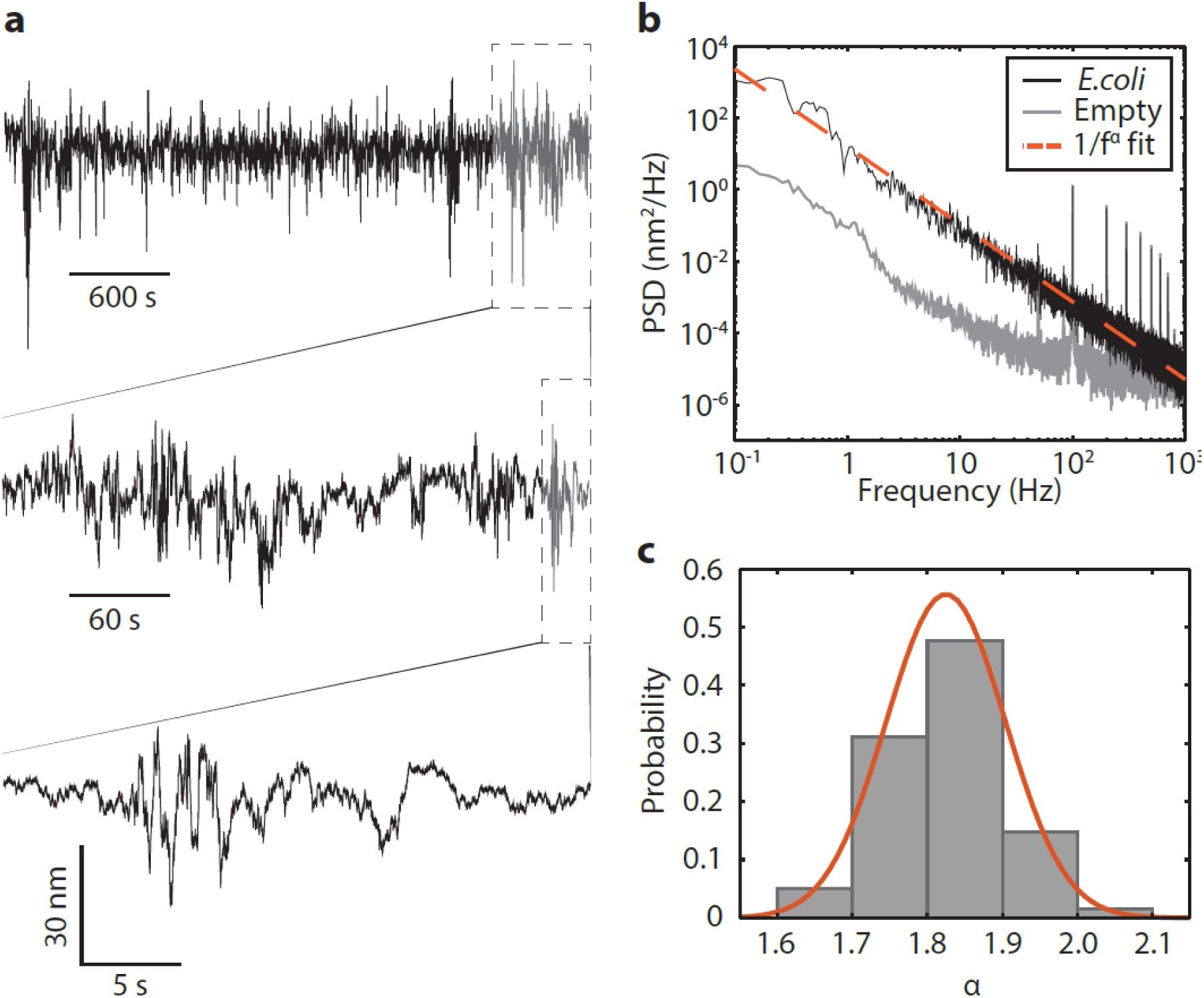
Motion of a single bacterium. a) Deflection *z*(t) versus time for a graphene drum with a single *E. coli* in LB, recorded for one hour. By zooming in on the part indicated in grey, while maintaining the same y-axis scale, it is observed that fluctuations are present over a wide range of timescales. b) Amplitude power spectral density (PSD) of the time-trace shown in 2a, of a live bacterium (black) and for the baseline from an empty drum (grey). Dashed orange line is a fit to 1/f^α^ spectrum with α=2.1. Peaks appear at harmonics of 50 Hz due to mains interference. c) Probability distribution of α from fitting 1/f^α^ noise. Orange line represents a Gaussian fit to the distribution, yielding an average value of α = 1.8 ± 0.1 (mean ± S.D.) (n= 277 samples).

## Impact of flagellar motility on nanomotion

While various origins of nanomotion have been proposed^3,4^, we speculate that flagellar motility constitutes the major source. To clarify its role on the bacterial forces generated, we compare the nanomotion of four *E. coli* strains (Figure 3a) that were genetically modified to have varying levels of motility: a hyper-motile strain with a larger number of flagella compared to wildtype, a minimally motile strain that lacks the regulatory IS1 element for the flagellum synthesis^21,22^, a non-motile strain with disabled flagellar motors, and a flagella-less strain where the motors are functional but flagella are lacking. As a fifth case, we studied the overall influence of ion pumps on the nanomotion by administering cadaverine, a drug, that blocks ionic transport through the cell membrane^23^ and thus reduces cell motility.

**Figure 3.**
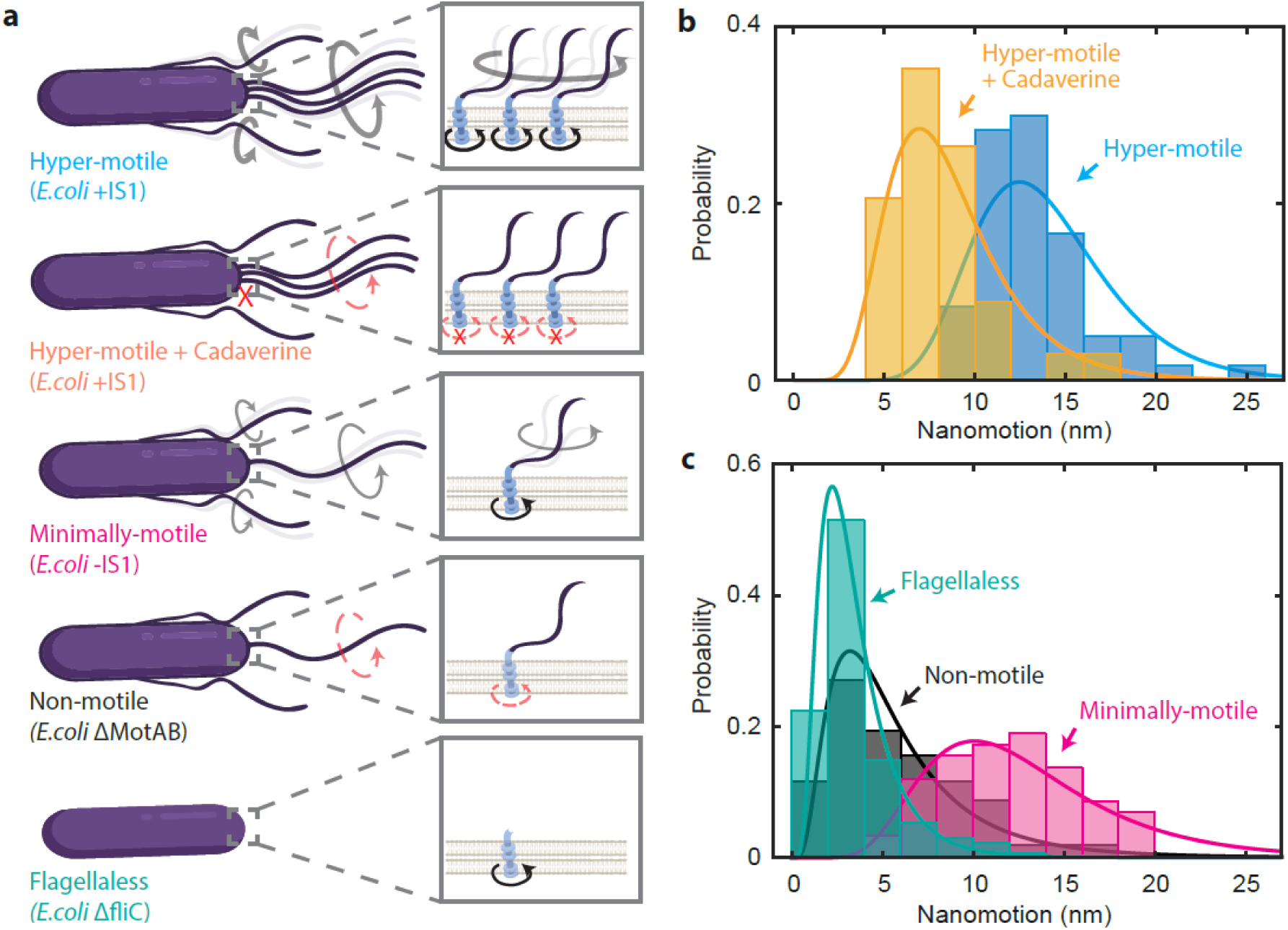
Impact of flagellar motility on nanomotion. a) To study the influence of motility on nanomotion, the following strains are compared: hypermotile, hypermotile with motility impaired by cadaverine, minimally motile by a genetic blocker, non-motile by gene deletion, and flagella-less. Red cross indicates a blocked ion pump and red dashed arrow indicates a disabled motor. b) Histogram of the motion amplitude σ of hyper-motile bacteria before (blue, n=60) and after (yellow, n=34) administering cadaverine, showing reduction of nanomotion. c) Motion σ of minimally-motile *E. coli* (purple, n=58) is compared to the non-motile strain (black, n=103) and the flagella-less strain (turquoise, n=169). The non-motile and flagella-less strains showed significantly lower motion than the minimally motile strain. Lines represent lognormal fits to the distributions.

The histograms in Figure 3b compare the motion of hyper-motile bacteria before and after exposure to cadaverine. The motion amplitude σ is observed to be substantially lowered after adding the drug (the median reduced from *σ* = 13.4 nm to 7.0 nm before and after administering cadaverine, respectively), indicating that the bacterial motion was strongly reduced, although it did not get fully quenched. The level of motility was observed to have a large influence on the magnitude of the nanomotion signal, as shown in Figure 3c. We observed that the nanomotion from the strains with both functional flagella and motors (median of *σ* = 13.4 nm for hyper-motile and *σ* = 12.6 nm for minimally-motile strains) was significantly larger than from strains in which either the motor was disabled or the flagella was removed (median variance *σ* = 5.3 nm for non-motile, and *σ* = 2.6 nm for flagella-less strains). We conclude that the observed differences in nanomotion are mainly induced by the activity of flagella, since the nanomotion disappeared in the flagella-less strain and the amplitude clearly correlates with the activity of the flagella.

## Antibiotic susceptibility tests on single bacteria

Subsequently, we explored if antibiotic susceptibility tests can be performed on single *E. coli* bacteria by monitoring nanomotion of graphene drums. To test the efficacy of different antibiotics, we measured the nanomotion variance *σ*^2^ of each drum for 30 seconds, both before and 1 hour after administering an antibiotic above its minimum inhibitory concentration (MIC). Figure 4a shows the 6 different antibiotics that we tested and their mode of action. For the antibiotics rifampicin, ciprofloxacin, DNP, and chloramphenicol, a decrease in the nanomotion was observed (Figure 4b-f and Table 1). Initially, a median motion amplitude *σ* = 7 nm is observed for the AB1157 *E. coli* strain, but quickly after administering the antibiotic the amplitudes drop to median values around *σ* = 3 nm. These results show that one can use graphene drums for testing antibiotic susceptibility based on nanomotion.

**Figure 4.**
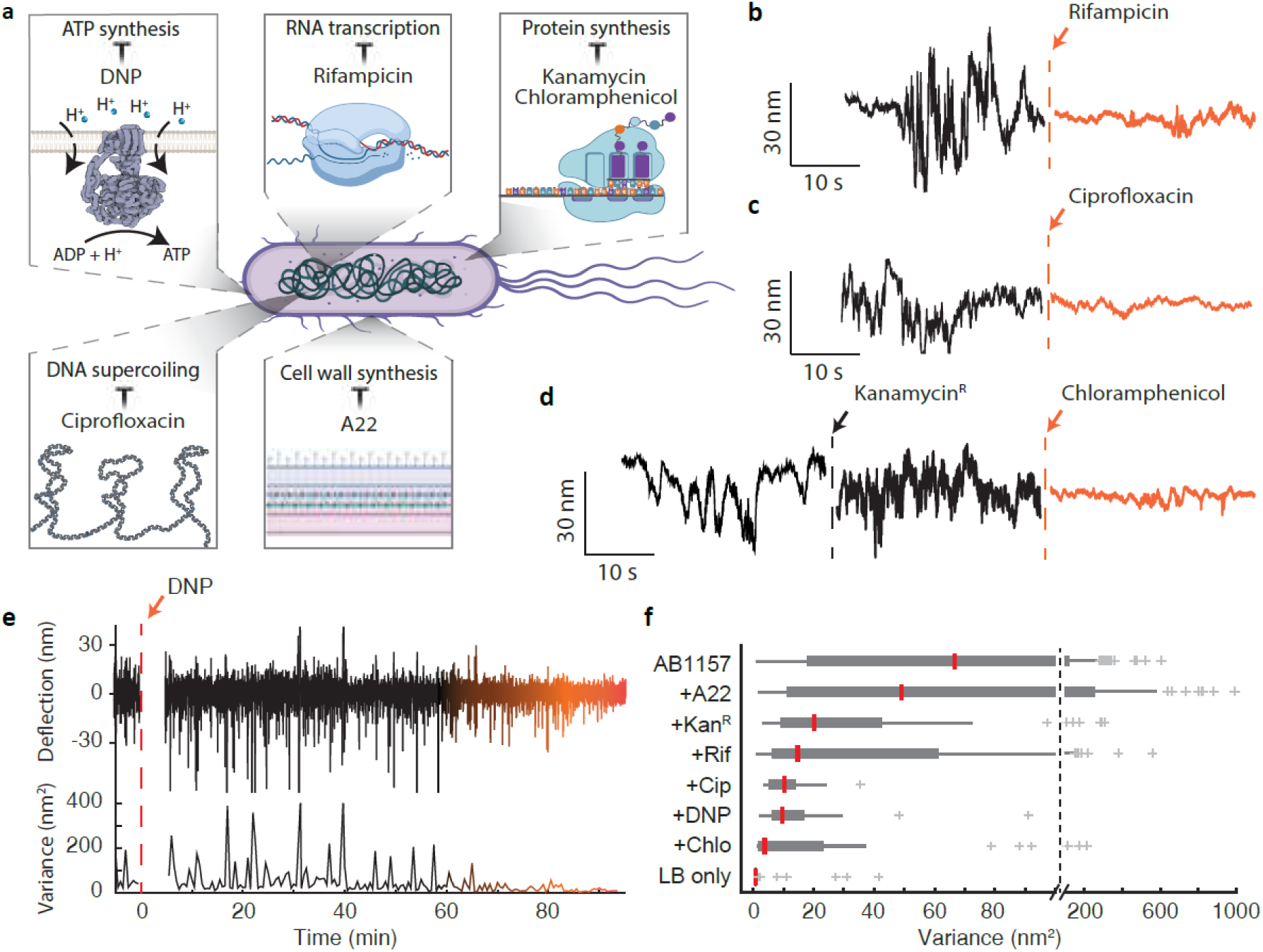
Single-cell antibiotic sensitivity screening using graphene. a) Schematic of antibiotics used in this work and their respective mode of action. b,c) Recorded motion of a drum with *E. coli* before (black) and after exposure to antibiotics (orange). d) Recorded motion of a drum with a kanamycin-resistant *E. coli* bacterium before (black), and after 1 hour of exposure to kanamycin, and subsequently after 1 hour of exposure to chloramphenicol. e) Recorded deflection (upper trace) that starts 6 minutes after DNP drug injection at t=0 min (vertical orange dashed line). A moving average is used to calculate the variance (below) using a window of 30’. f) Box plot for all measurements after administering the corresponding drug to the *E. coli* strain AB1157 (n=277), +A22 (n=108), +kanamycin (n=33), +rifampicin (n=83), +ciprofloxacin (n=36), +DNP (n=27), +chloramphenicol (n=33), as well as for empty control drums in LB (n=80). Box plot indicates the 25th, 50th (red line is the median) and 75th percentiles, whereas whiskers extend to maximum 1.5 times the interquartile distance. Outliers are indicated by a cross.

**Table 1.**
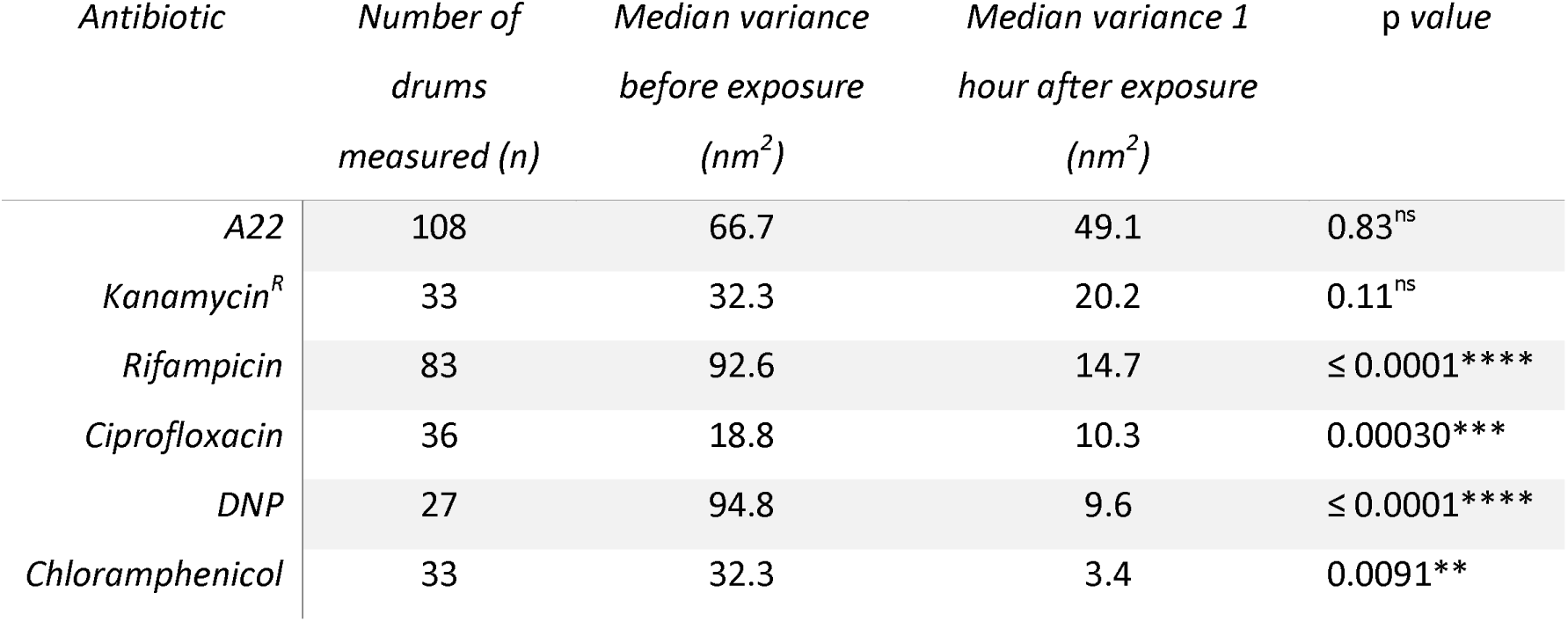
Efficacy of antibiotics measured 1 hour after exposure. Median value of the variance before and after exposure are compared, and the probability (p-value) that the drug has no effect on the nanomotion variance is evaluated using a two-tailed rank test. For all antibiotics except A22, a two-tailed Wilcoxon signed rank test is performed for paired measurements before and after exposure to the antibiotic. For A22, a two-tailed Wilcoxon rank sum test is performed with respect to *E. coli* AB1157. Superscript R indicates antibiotic resistance and significance is expressed using the asterisk convention.

To test whether graphene drums are able to distinguish resistant cells, we used *E. coli* cells with chromosomal KanR resistance gene^24^. When these cells were exposed to kanamycin, we observed no change in the motion amplitude (*σ* ∼ 5 nm) (Figure 4d). However, when we subsequently exposed the same cells to chloramphenicol, we did observe a decrease in the signal with respect to the initial nanomotion (down to *σ*= 1.8 nm). Additionally, we treated *E. coli* cells with A22 which stalls cell wall synthesis (see Supplementary Note 4). We used sub-MIC concentrations of the drug, in order to test if the cell-wall synthesis was a contributing factor to the observed vibrations, without killing the cells. In this case, the variance was found to be similar to that of the untreated cells, even though the alteration of the cell wall was visible under an optical microscope as the bacteria were clearly rounded and lost their typical rod shape^25-27^. In contrast to the effect of the other antibiotics, disrupting the cell-wall synthesis was not observed to result in a reduction in the nanomotion.

Besides detecting differences in nanomotion between strains, or after administering antibiotics, the graphene platform also offers the possibility of real-time probing of the decrease in vibration amplitude, providing on-the-fly information on the route to bacterial death. From long-time trace measurements such as Figure 4e (and Supplementary Note 6), we found that most of the nanomotion fades within the first hour after exposure to antibiotics. This experiment demonstrates the potential of graphene devices as an indicator of bacterial physiology, and opens new routes for determining the temporal response of bacteria to antibiotics at single cell level.

## Discussion

We present an ultrasensitive platform that uses graphene drums to measure nanomotion of single bacterial cells. Single *E. coli* bacteria were observed to produce peak fluctuations of up to 60 nm in amplitude, that correspond to forces of up to 6 nN as inferred from the graphene membrane stiffness of *k* ≈ 0.1 N/m (see Methods). These forces are larger than the typical forces generated by a single molecular motor^28^ (*F* ∼ 10 pN) or a single flagellar motor^29,30^ (*F* ∼ 100 pN), but are similar to forces measured by AFM spectroscopy on membranes of single *Saccharomyces cerevisiae* cells^31^ (*F* ∼ 10 nN). By comparing the nanomotion of different strains of bacteria, we conclude that flagellar motion is the major contributing factor to the nanoscale vibrations. It is worth noting though, that flagellar motility is not the only source of nanomotion, as it was observed even in flagella-less *E. coli*, albeit at significantly lower amplitude.

Recent reports call for the development of effective diagnostic tools to detect antimicrobial resistance and slow down the emergence of multi-drug resistant bacteria by prescribing the correct drug^32^. Our antibiotic susceptibility experiments demonstrated that the graphene drum sensing platform can trace the effect of antibiotics on bacterial nanomotion in real-time. This opens the way to fast, label-free susceptibility testing down to the single bacterial level. In comparison to other techniques for detecting antibiotic susceptibility^33^, the method presented here stands out in terms of sensitivity and speed, offering for the first time the capability to quantify the nanomotion at the level of individual bacteria within a timeframe of 30 seconds. The small size of the graphene drums enables massive parallelization, allowing, in principle, millions of cells to be monitored in parallel in the presence of antibiotics. This would facilitate the identification of important minority fractions like persister cells, that are related to the emergence of antibiotic resistance^34^. Similar benefits might apply in the field of personalised medicine, where the right antibiotic can be rapidly selected based on the nanomotion response.

Furthermore, directed evolution experiments may benefit from this technique as a fast selection and screening method^35^, as the density of over 10.000 nanomotion sensors/mm^2^ can result in a greatly increased throughput as compared to 96-well plates or petri-dish culturing. With the significant reduction in size and increase in sensitivity presented in this work, nanomotion detection potentially can evolve into an important non-invasive monitoring tool in cell biology and provide new routes for rapid screening tests in personalized medicine and drug development.

## Methods

### Bacterial strains

For antibiotics susceptibility experiments, FW2179, a derivative of *E. coli* AB1157 strain, described previously in ^27^, was used. Hypermotile (MG1655(+IS1)), minimally motile (MG1655(-S1)), non-motile (MG1655ΔmotAB) and flagella-less (MG1655ΔfliC) strains that were described previously in ^36^, were a kind gift from Dr. Bertus Beaumont from TU Delft.

### Sample preparation

For experiments with *E. coli* cells, we grew cells in LB media overnight at 30°C to reach the late exponential phase. On the day of the experiment, the overnight culture was refreshed (1:100 volume) for 2.5 hours on fresh LB medium at 30°C to reach OD_600_=0.2-0.3. Then 1 ml of the refreshed culture was mixed with APTES (Sigma-Aldrich) to reach a final concentration of 0.1% APTES (volumetric). This acts as a binder between the bacteria and the chips^37^. A cuvette with a graphene covered chip inside was then filled with the solution. The chamber was left for 15 minutes in a horizontal position to deposit the bacteria on the surface. Afterwards, the chamber was placed in an upright position to prevent additional bacteria from depositing and maintain an average coverage of a single bacterium per drum. An optical microscope (Keyence VHX-7000) was used to inspect the sample. The cuvette was then placed in the optical nanomotion detection setup (Figure 1a). The setup was equipped with nano positioners (Attocube ECSx5050) that allow for automated scanning over an array of drums. The motion of the bacterium is transduced on the drum and recorded using a digital oscilloscope (Rohde & Schwarz RTB2004). For each drum a trace was recorded for at least 30’ with a sampling rate of at least 500 Hz. The measurements were performed in an air-conditioned room with a temperature of 21 degrees Celsius. After measuring the sample for one hour and collecting approximately 60 time-traces of different drums, antibiotics were added to the solution at the concentration given in Table 2. The antibiotic was left to work for one hour (unless otherwise stated) and afterwards a new round of measurements was performed on the same array of graphene drums.

**Table 2:** Table describing the types and concentrations of antibiotics used (see also Figure 4a).

| *Antibiotic* | *Target* | *Mechanism* | *Concentration* |
| --- | --- | --- | --- |
| *A22* | Cell wall synthesis | Inhibits MreB filament polymerization | 5 $\mu\text{g/ml}$ |
| *Kanamycin* | Translation | Binds ribosome and interferes in elongation of polypeptide chain elongation | 50 $\mu\text{g/ml}$ |
| *Chloramphenicol* | Translation | Binds to ribosome and inhibits binding of tRNA | 34 $\mu\text{g/ml}$ |
| *Ciprofloxacin* | DNA Supercoiling homeostasis | Traps topoisomerase and DNA in a complex, inhibits DNA rejoining after cleavage | 15 $\mu\text{g/ml}$ |
| *Cadaverine* | Ion transport | Induces closure of porins and inhibits ion transport over the membrane | 50 mM |
| *Rifampicin* | Transcription | Binds to RNAP and blocks the elongating RNA molecule | 50 $\mu\text{g/ml}$ |
| *DNP* | $\text{H}^+$ gradient across the membrane | Inhibits ATP synthesis | 2 mM |

### Substrates

The substrates were 5×5 mm^2^ silicon chips with a 285 nm layer of silicon oxide that were patterned with circular holes by a reactive ion etch, where the silicon acts as a stop layer. Chemical Vapour Deposited (CVD) bilayer graphene was supplied, transferred, and suspended over the circular holes by Graphenea with a dry transfer method. The quality of the graphene drums was inspected by Scanning Electron Microscopy (SEM) and optical microscopy. Suspended circular drums with a diameter of 8 µm were used for the experiments.

### Antibiotics

The antibiotics used in this work are listed in table 2.

### Amplitude calibration

Here we describe how the drum deflection *z*(t) was obtained from the reflected intensity variations *I*(t) of the red laser that was reflected by the photodiode voltage *V*_*pd*_(t). We first define the reflection coefficient *R*(t)=*I*(t)/*I*_*0*_, where *I*_*0*_ is the incident light intensity and *I*(t) is the reflected light intensity. The reflection coefficient *R*(t) depends on the optical characteristics of the cavity formed between the graphene and the silicon and the position *z*(t) of the graphene membrane. Light passes subsequently through three media with the following refractive indices: LB media with *n*_LB_ = 1.34 – 0.0007i, graphene with *n*_gr_ = 2.7-1.6i, air with *n*_air_ = 1, and finally the light was reflected from the silicon mirror *n*_si_ = 4.2-0.06i, where i is the imaginary unit. Together, the semi-transparent graphene layer and the reflective silicon form a Fabry-Pérot cavity. The reflected light is modulated by the graphene drum moving through the optical field, and the reflection coefficient *R* = *I*/*I*_*0*_ can be described by the following equation^38^

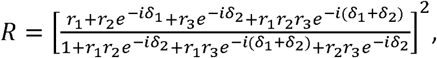

where —, —and —, and the exponent is the phase difference that the light of wavelength acquires while travelling through a medium of thickness. In this case —and —, with *t*_*air*_=*g* + *z*(t). The reflectivity of the cavity depends on the number of graphene layers and the cavity depth, as plotted in Extended Data Figure 1a, where the reflectivity for bilayer graphene is indicated by a red line. The design cavity depth is 285 nm, however the drums bulged down by typically 60 nm under pressure of the liquid as can be seen in the liquid AFM image (Supplementary Note 5). Therefore, we consider that the effective cavity depth was *g*=225 nm. Then, we normalized the reflectivity by dividing it over *R* at a cavity depth of 225 nm (*R*_*0*_), to find the slope around that point, which equals *φ*=d(*R*(t)/*R*_*0*_)/dz=*−0*.*0038 nm*^-1^, as indicated in Extended Data Figure 1b.

Data was gathered by an oscilloscope measuring the voltage *V*_*pd*_(t) from the photodiode that is proportional to the reflected light intensity and is operated in its linear range. The gathered time trace was normalized by division over its average, *V*_*norm*_=*V*_*pd*_(t)/<*V*_*pd*_(t)>, and a linear fit was subtracted from the data to eliminate the effects of drift during the measurement. Using the calibration factor *φ*, the deflection *z*(t) was calculated as *z*(t)= *[V*_*pd*_(t)/<*V*_*pd*_(t)> − *1]*/ *φ*.

**Extended Data Figure 1:**
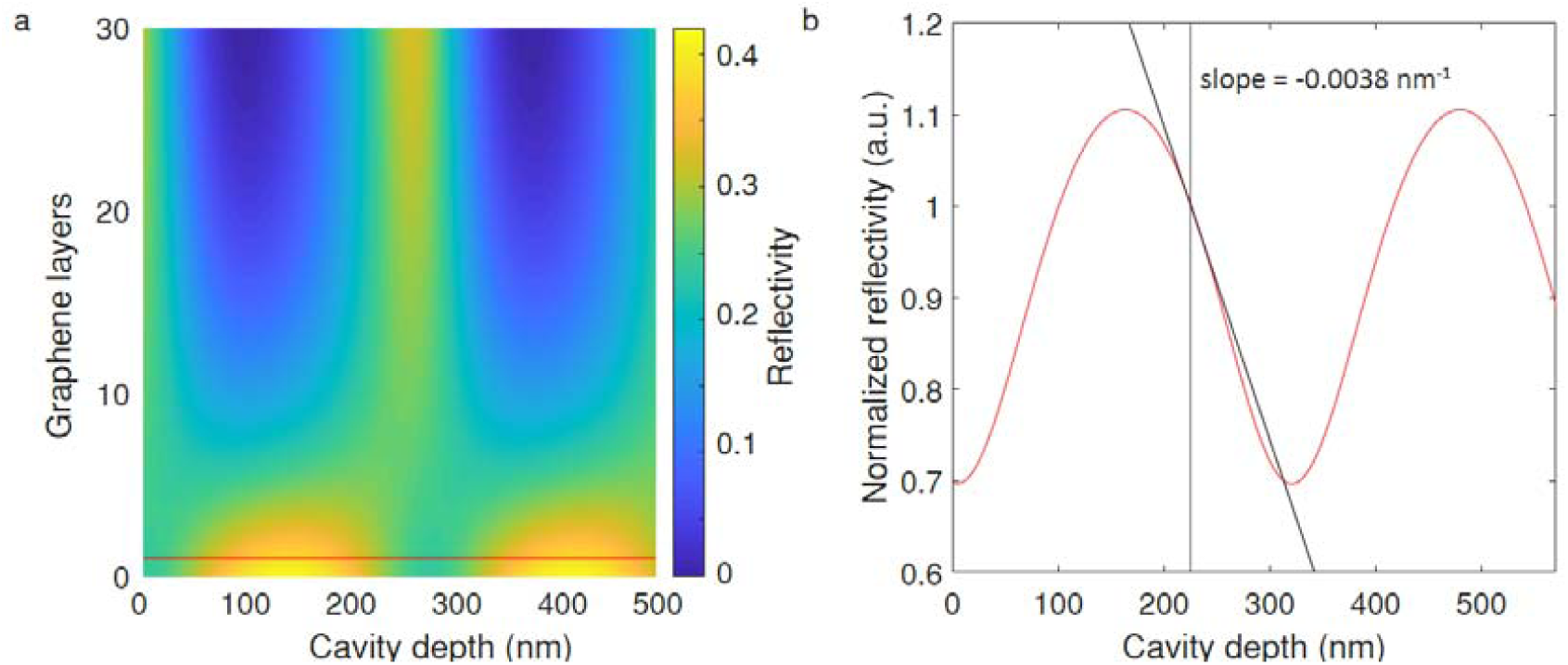
a) Reflectivity as a function of the cavity depth *t*_*air*_ and the number of graphene layers. b) Normalized reflectivity versus cavity depth *t*_*air*_.

While the current nanomotion detection technique works well for qualitative analysis of changes in the bacterial nanomotion in time, there are several approximations made in the conversion from the nanomotion-induced light-intensity variations detected by the photodiode to a nanomotion amplitude in nm. First of all, the nanomotion generated by a bacterium may depend on its position on the drum, which could cause experimental variations. In our calculations of the force, we assume that a single bacterium is centered on the drum. Moreover, in the optical model, the cavity underneath the graphene is assumed to be filled by air. The use of bilayer graphene minimizes the chances that small defects cause leakage and liquid AFM measurements (see Supplementary Note 5) also showed that the graphene membranes bulge down, which is to be expected if the cavity is air filled. Finally, the bacterium is attached to the surface of the graphene and is likely to be in the laser beam path. The refractive index of an *E. coli* bacterium^39,40^ (n = 1.33) is very close to that of the LB medium (n = 1.34), causing the bacteria to be nearly transparent and therefore we estimate this to have negligible impact on the nanomotion amplitude determination.

### Estimation of the stiffness of a graphene drum

We estimate the stiffness *k*_*1*_ of the circular graphene drum with area *A* = *50* µm^2^ based on the deflection *z* at the centre of the membrane with respect to a flat configuration induced by uniform liquid pressure *P* in the cuvette. Hooke’s law prescribes that the stiffness can be found by equating forces:

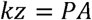

The graphene drum is immersed 1 cm below the surface of the liquid and is therefore under a uniform pressure of 100 Pa. Under these conditions the graphene is found to deflect 60 nm downward, as measured by liquid AFM (see supplementary note 5). By inserting these values in the equation above, we find *k* = 0.14 N/m. Our estimate of the stiffness of graphene drums corresponds to values reported in literature^41-43^, which typically range from 0.05 to 1 N/m.

### Statistics

Since the data reported in the manuscript are not normally distributed, we relied on non-parametric tests for statistics. We represent the median and quartiles of data in boxplots, in accordance with the use of non-parametric tests. We use a signed rank test whenever repeated measurements on the same drum are available (i.e. antibiotic susceptibility test), and rank sum test for comparison between strains. We used Matlab built-in functions for statistical analysis. All statistical tests were two-sided. On all figures, the following conventions are used: ns: 0.05 < p, *: 0.01 < p < 0.05, **: 0.001 < p < 0.01, ***: 0.0001 < p < 0.001, ****: p < 0.0001. We report a significant difference in results if p < 0.01.

## Data availability

The datasets of this study are available from the corresponding author on request.

## Acknowledgements

Financial support was provided by from the European Union’s Horizon 2020 research and innovation programme under ERC starting grant ENIGMA (802093), ERC PoC GRAPHFITI (966720), Graphene Flagship (785219 and 881603), and the ERC Advanced Grant LoopingDNA (883684), as well as the Netherlands Organisation for Scientific Research (NWO/OCW), as part of the NanoFront and BaSyC programs, and by the Swiss National Science Foundation (P300P2_177768). We acknowledge Graphenea for providing and transferring the bilayer graphene used in this research. We thank Miloš Tišma for help in the wetlab, Bertus Beaumont and Alexis Derumigny for discussions, and Tim de Visser, Samir Mohammad, Nick Wansink, and Sebastiaan van Putten for their contributions to the nanomotion detection setup. Schematics in Figure 3a and 4a were created with Biorender.com.

## Author contributions

A.J. and F.A. conceived the idea. I.E.R. and A.J. collected the data. I.E.R. constructed the setup and performed the interferometry experiments. A.J. performed the bacterial manipulation. All authors designed the experiments. The project was supervised by C.D., P.G.S. and F.A. All authors contributed to the data analysis, interpretation of the results and writing of the manuscript.

## References

1 Glass, L. & Mackey, M. C. From clocks to chaos: the rhythms of life. XVII, 248 p. : illustrations ; 4 cm.(Princeton University Press, 1988).

2 Longo, G. et al. Rapid detection of bacterial resistance to antibiotics using AFM cantilevers as nanomechanical sensors. Nat Nanotechnol 8, 522–526 (2013).

3 Kohler, A. C., Venturelli, L., Longo, G., Dietler, G. & Kasas, S. Nanomotion detection based on atomic force microscopy cantilevers. The Cell Surface 5 (2019).

4 Venturelli, L. et al. A perspective view on the nanomotion detection of living organisms and its features. J Mol Recognit, e2849 (2020).

5 Jülicher, F. Mechanical oscillations at the cellular scale. Comptes Rendus de l’Académie des Sciences-Series IV-Physics-Astrophysics 2, 849–860 (2001).

6 Ruggeri, F. S. et al. Amyloid single-cell cytotoxicity assays by nanomotion detection. Cell Death Discovery 3, 17053 (2017).

7 Willaert, R. G. et al. Single yeast cell nanomotions correlate with cellular activity. Science Advances 6, eaba3139 (2020).

8 Kohler, A.-C. et al. Yeast Nanometric Scale Oscillations Highlights Fibronectin Induced Changes in C. albicans. Fermentation 6, 28 (2020).

9 Cadart, C., Venkova, L., Recho, P., Lagomarsino, M. C. & Piel, M. The physics of cell-size regulation across timescales. Nature Physics 15, 993–1004 (2019).

10 Li, M., Xi, N., Wang, Y. & Liu, L. Advances in atomic force microscopy for single-cell analysis. Nano Research 12, 703–718 (2019).

11 Martínez-Martín, D. et al. Inertial picobalance reveals fast mass fluctuations in mammalian cells. Nature 550, 500–505 (2017).

12 Krieg, M. et al. Atomic force microscopy-based mechanobiology. Nature Reviews Physics 1, 41–57 (2019).

13 Arbore, C., Perego, L., Sergides, M. & Capitanio, M. Probing force in living cells with optical tweezers: from single-molecule mechanics to cell mechanotransduction. Biophysical reviews 11, 765–782 (2019).

14 Otto, O. et al. Real-time deformability cytometry: on-the-fly cell mechanical phenotyping. Nature Methods 12, 199–202 (2015).

15 Bennett, I., Pyne, A. L. B. & McKendry, R. A. Cantilever Sensors for Rapid Optical Antimicrobial Sensitivity Testing. ACS Sensors 5, 3133–3139 (2020).

16 Syal, K. et al. Antimicrobial Susceptibility Test with Plasmonic Imaging and Tracking of Single Bacterial Motions on Nanometer Scale. ACS Nano 10, 845–852 (2016).

17 Steeneken, P. G., Dolleman, R. J., Davidovikj, D., Alijani, F. & van der Zant, H. S. Dynamics of 2D Material Membranes. 2D Materials 8 (2021).

18 Rosłoń, I. E. et al. High-frequency gas effusion through nanopores in suspended graphene. Nature Communications 11, 6025 (2020).

19 Davidovikj, D. et al. Visualizing the Motion of Graphene Nanodrums. Nano Letters 16, 2768–2773 (2016).

20 Lissandrello, C. et al. Nanomechanical motion of Escherichia coli adhered to a surface. Appl Phys Lett 105, 113701 (2014).

21 Barker, C. S., Prüß, B. M. & Matsumura, P. Increased motility of Escherichia coli by insertion sequence element integration into the regulatory region of the flhD operon. Journal of bacteriology 186, 7529–7537 (2004).

22 Leatham, M. P. et al. Mouse Intestine Selects Nonmotile flhDC Mutants of Escherichia coli MG1655 with Increased Colonizing Ability and Better Utilization of Carbon Sources. Infection and Immunity 73, 8039–8049 (2005).

23 Sowa, Y. et al. Direct observation of steps in rotation of the bacterial flagellar motor. Nature 437, 916–919 (2005).

24 Bremer, E., Silhavy, T. J. & Weinstock, G. M. Transposable lambda placMu bacteriophages for creating lacZ operon fusions and kanamycin resistance insertions in Escherichia coli. Journal of Bacteriology 162, 1092–1099 (1985).

25 Karczmarek, A. et al. DNA and origin region segregation are not affected by the transition from rod to sphere after inhibition of Escherichia coli MreB by A22. Molecular microbiology 65, 51–63 (2007).

26 Japaridze, A., Gogou, C., Kerssemakers, J. W., Nguyen, H. M. & Dekker, C. Direct observation of independently moving replisomes in Escherichia coli. Nature communications 11, 1–10 (2020).

27 Wu, F. et al. Direct imaging of the circular chromosome in a live bacterium. Nature Communications 10, 2194 (2019).

28 Visscher, K., Schnitzer, M. J. & Block, S. M. Single kinesin molecules studied with a molecular force clamp. Nature 400, 184–189 (1999).

29 Ryu, W. S., Berry, R. M. & Berg, H. C. Torque-generating units of the flagellar motor of Escherichia coli have a high duty ratio. Nature 403, 444–447 (2000).

30 Mandadapu, K. K., Nirody, J. A., Berry, R. M. & Oster, G. Mechanics of torque generation in the bacterial flagellar motor. Proceedings of the National Academy of Sciences 112, E4381–E4389 (2015).

31 Pelling, A. E., Sehati, S., Gralla, E. B., Valentine, J. S. & Gimzewski, J. K. Local Nanomechanical Motion of the Cell Wall of Saccharomyces cerevisiae. Science 305, 1147–1150 (2004).

32 WHO. No Time to Wait: Securing the future from drug-resistant infections. (2019).

33 Van Belkum, A. et al. Innovative and rapid antimicrobial susceptibility testing systems. Nature Reviews Microbiology 18, 299–311 (2020).

34 Barrett, T. C., Mok, W. W., Murawski, A. M. & Brynildsen, M. P. Enhanced antibiotic resistance development from fluoroquinolone persisters after a single exposure to antibiotic. Nature communications 10, 1–11 (2019).

35 Cobb, R. E., Chao, R. & Zhao, H. Directed evolution: Past, present, and future. AIChE Journal 59, 1432–1440 (2013).

36 Gauger, E. J. et al. Role of Motility and the flhDC Operon in Escherichia coli MG1655 Colonization of the Mouse Intestine. Infection and Immunity 75, 3315–3324 (2007).

37 Louise Meyer, R. et al. Immobilisation of living bacteria for AFM imaging under physiological conditions. Ultramicroscopy 110, 1349–1357 (2010).

38 Blake, P. et al. Making graphene visible. Applied Physics Letters 91, 063124 (2007).

39 Balaev, A., Dvoretski, K. & Doubrovski, V. Refractive index of escherichia coli cells. Vol. 4707 SFM (SPIE, 2002).

40 Liu, P. Y. et al. Real-time Measurement of Single Bacterium’s Refractive Index Using Optofluidic Immersion Refractometry. Procedia Engineering 87, 356–359 (2014).

41 Suk, J. W., Piner, R. D., An, J. & Ruoff, R. S. Mechanical Properties of Monolayer Graphene Oxide. ACS Nano 4, 6557–6564 (2010).

42 Davidovikj, D. et al. Nonlinear dynamic characterization of two-dimensional materials. Nature Communications 8 (2017).

43 Castellanos-Gomez, A., Singh, V., van der Zant, H. S. & Steele, G. A. Mechanics of freely-suspended ultrathin layered materials. Annalen der Physik 527, 27–44 (2015).

