## Supplemental Notes 1-6 for "Probing nanomotion of single bacteria with graphene drums"

### SUPPLEMENTARY INFORMATION:
### Probing nanomotion of single bacteria with graphene drums

I.E. Rosłoń, A. Japaridze, P.G. Steeneken, C. Dekker, and F. Alijani*
*Delft University of Technology, Delft, The Netherlands*

#### CONTENTS

Supplementary Note 1: Optical characterization 2

Supplementary Note 2: Control experiments on drums without bacteria 3

Supplementary Note 3: Extended measurement data - strains 3

Supplementary Note 4: Extended measurement data - antibiotics 5

Supplementary Note 5: Liquid AFM measurement 7

Supplementary Note 6: Measurement on exfoliated graphene 8

References 8

*

#### SUPPLEMENTARY NOTE 1: OPTICAL CHARACTERIZATION

Chips with graphene drums, dimensions 5x5 $mm^2$, were first inspected under a microscope. Not all drums were successfully suspended after fabrication of the device and transfer of the graphene. The drums which were intact could be recognized as they showed up darker than collapsed drums under a blue filter, as can be seen in Supplementary Figure 1a. Once the position of intact drums in the array was known, the chip was fixed inside a cuvette and the growth medium containing bacteria was added. The chip is placed in horizontal position, such that the bacteria sediment on the surface of the chip. Approximately 20 minutes of deposition was required to obtain an average coverage of 1 bacterium per drum in our setup with a growth medium at $OD_{600} = 0.2 - 0.3$. Once this time had elapsed, the chip was placed vertically to stop deposition of more bacteria. The sample was again inspected under the microscope to check if the bacteria are well adhered. Supplementary Figure 1b shows an example of an array of graphene drums with adhered cells. The sample was then placed in the laser interferometry setup in which the nanomotion can be observed by focusing the laser spot on one of the drums.

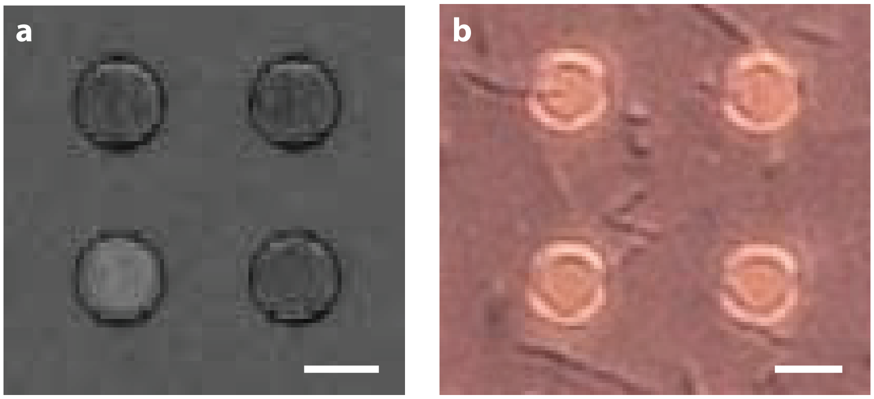


SUPPLEMENTARY FIG. 1. The array of graphene drums was inspected optically before and after deposition of the bacteria. a) Intact drums can be recognized as they show up darker than collapsed drums under a blue filter. Only the intact drums were measured. b) An array of membranes with *E.coli* after deposition. The chip was placed in upright position within the liquid chamber, to ensure that the bacteria are well attached and no more bacteria attach on the surface. Scalebars are 10 $\mu$m long.

#### SUPPLEMENTARY NOTE 2: CONTROL EXPERIMENTS ON DRUMS WITHOUT BACTERIA

We performed control experiments to establish whether the bacteria are the source of the nanomotion. First, a experiment was performed without bacteria on an array of suspended drums in LB growth medium. We detected a motion amplitude below $\sigma^2 = 4$ nm on these drums, which is the noise floor for our measurements and is referred to in the main text as baseline. Three typical traces are shown in Supplementary Figure 2.

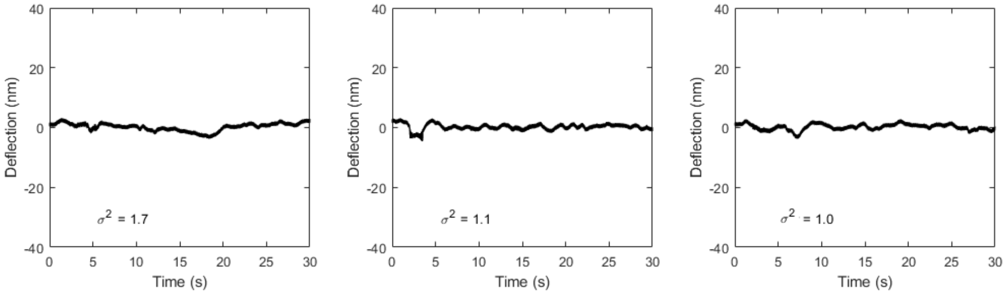


SUPPLEMENTARY FIG. 2. Measurement of nanomotion on drums without bacteria in LB.

We also performed a second type of control experiments to establish that the graphene drums are needed to transduce and read the nanomotion. In this experiment the laser was focused on AB1157 bacteria attached to the $Si/SiO_2$ substrate, outside of the suspended area. Here, we also detected a motion amplitude below $\sigma^2 = 4$ nm, much lower than the signals from bacteria on suspended graphene. Three typical traces are shown in Supplementary Figure 3.

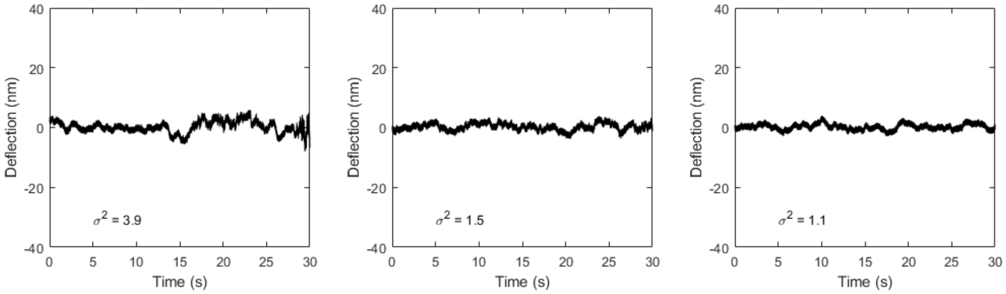


SUPPLEMENTARY FIG. 3. Measurement of nanomotion measured with the laser focused on a bacterium on the substrate, outside of the suspended area.

#### SUPPLEMENTARY NOTE 3: EXTENDED MEASUREMENT DATA - STRAINS

To study the influence of motility on the observed nanomotion, we studied five cases: hypermotile, hyper-motile with motility impaired by cadaverine, minimally motile by genetic blocker, non-motile by gene deletion, and flagellaless. Here, we show typical traces for each of these cases as a supplement to the data presented in Figure 3 in the main text.

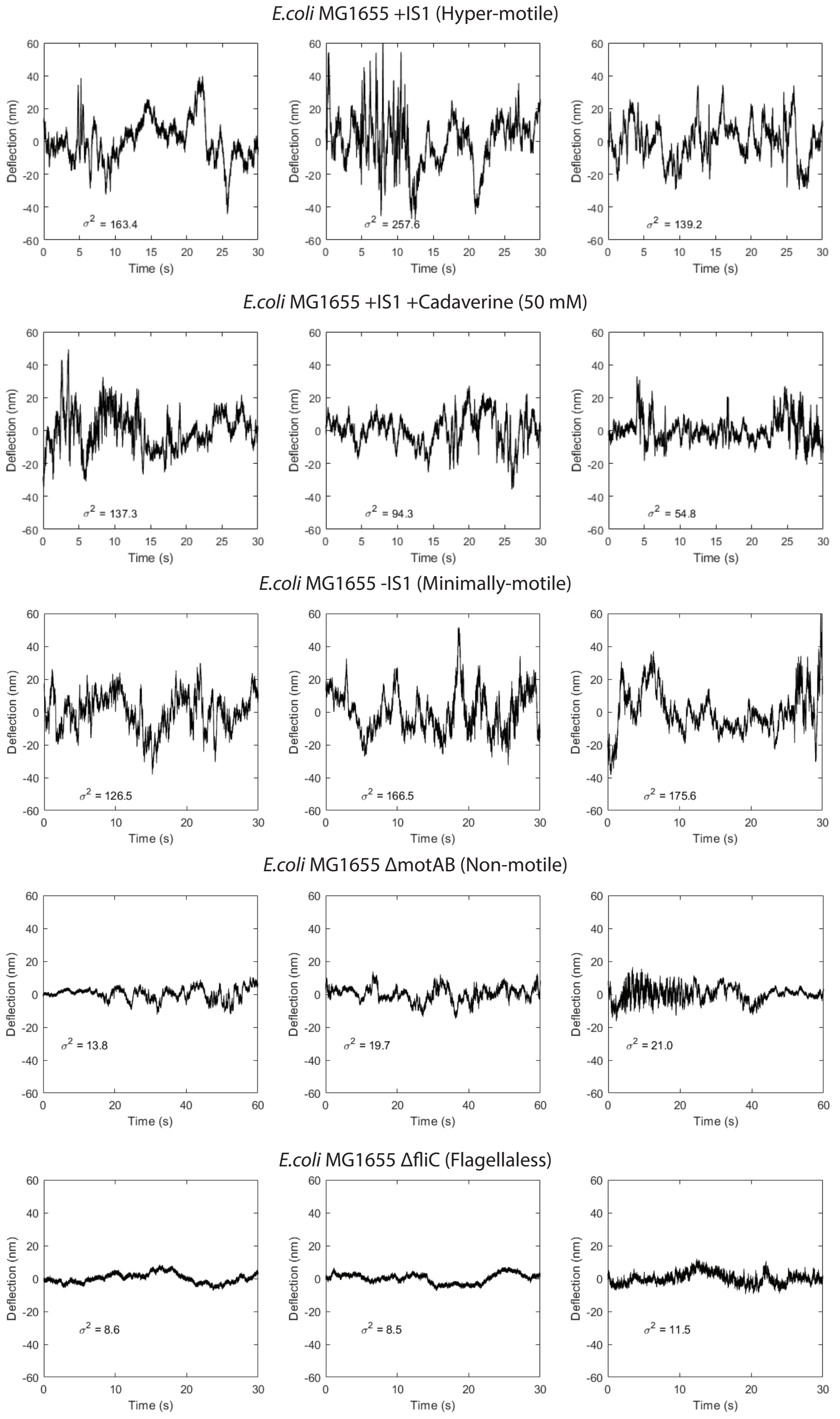


SUPPLEMENTARY FIG. 4. Impact of motility on observed nanomotion, extended graphs to Figure 3 in the main text. Each graph shows a measurements performed on single *E. coli*.

The various strains used in this work are listed in Table I. The median value of the nanomotion measured varied between strains. The motility of a strain was inferred from the biofilm formation on an agar plate [1], and is listed in the column *Motility* for comparison. The motility correlates with the magnitude of nanomotion we found in this study. This is consistent with the notion that motility is indeed an important contributor to the observed nanomotion.

| *Strain* | *Description* | *Motility* | *Median variance (nm$^2$)* | *Number of samples (-)* |
|---|---|---|---|---|
| AB1157 | Normal | Low | 67 | 277 |
| MG1655 (+IS1) | Hyper-motile | High | 170 | 60 |
| MG1655 (-IS1) | Minimally-motile | Medium | 158 | 58 |
| MG1655 (motAB) | Non-motile | None | 28 | 103 |
| MG1655 (fliC) | Flagellaless | None | 8 | 169 |

TABLE I. List of bacterial strains used in this research and their description.

#### SUPPLEMENTARY NOTE 4: EXTENDED MEASUREMENT DATA - ANTIBIOTICS

Here, time traces are presented for *E. coli* treated with A22. The variance was found to be similar to that of the untreated cells (Supplementary Figure 5). This means that disrupting cell wall synthesis does not kill the cell, and unlike all the other physiological processes that were blocked by the antibiotics it did not result in a significant change in the nanomotion. We note that in the case of A22, the cells were grown in the presence of the antibiotic before deposition on the graphene, and the alteration of the cell wall synthesis was first confirmed by optical microscopy as the bacteria are clearly rounded and have lost their typical rodshape.

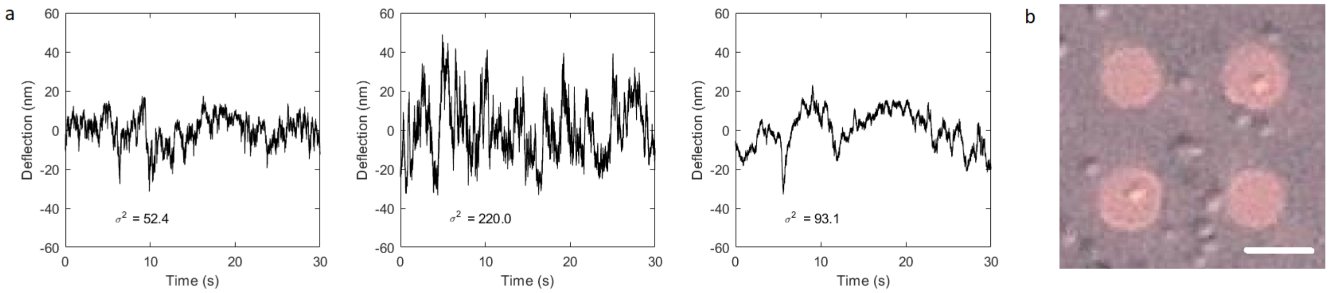


SUPPLEMENTARY FIG. 5. Impact of cell wall synthesis on observed nanomotion, extended graphs to Figure 4 in the main text. a) Each graph shows a measurements performed on *E. coli* pre-treated with A22. Nanomotion is similar to that of untreated *E. coli* AB1157, which indicates that the antibiotic did not kill the bacterium. Moreover, it tells that correct cell wall synthesis is not required to record nanomotion. b) Optical microscope imaging shows that the *E. coli* has lost its typical rod-shape after treatment with A22 and is now round. Scalebar is 10 $\mu$m.

In this supplement we also include extended data to Figure 4 in the main text, showing three more nanomotion traces for each of the antibiotics used on *E. coli* AB1157. Each graph in Supplementary Figure 6 represents two measurements performed on the same drum, before and after administering antibiotics. It is evident that for antibiotics to which *E. coli* is sensitive, the amplitude of oscillations decreases after exposition to the drug.

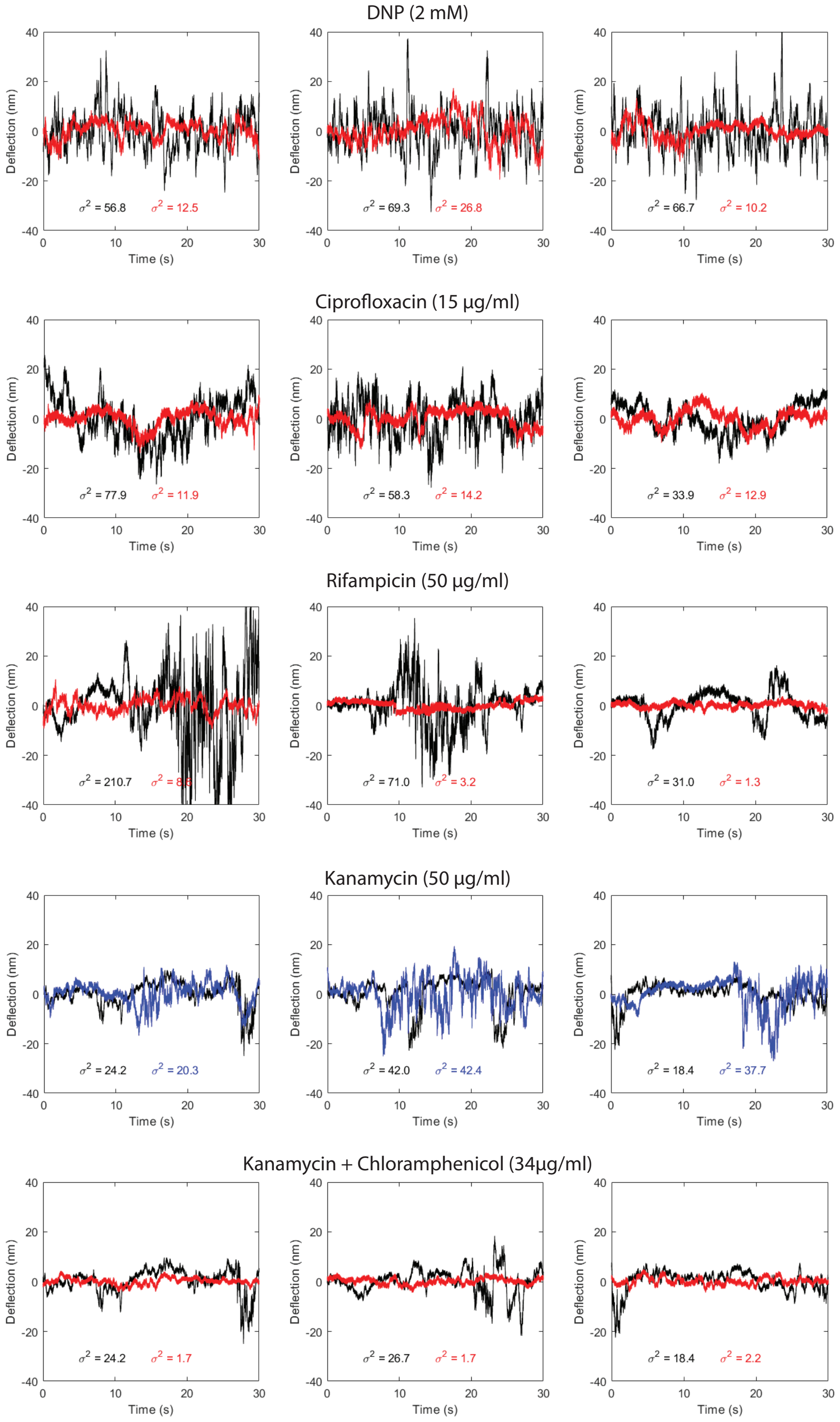


SUPPLEMENTARY FIG. 6. Single cell antibiotic sensitivity testing using graphene. Each graph shows two measurements performed on the same drum on *E. coli*, initially (black), and 1 hour after administering antibiotic (colored red and blue). The red traces show much lower signal, whereas blue traces do not show significant difference with respect to the original trace.

#### SUPPLEMENTARY NOTE 5: LIQUID AFM MEASUREMENT

The graphene drums were also characterized in liquid using atomic force microscopy (AFM). As shown in Supplementary Figure 7, the drums were intact and were bulging down by pressure. We find that the drums deflect by typically 35 to 60 nm, depending on the sample. Supplementary Figure 8 shows a bacterium adhered to the surface of the suspended graphene drum.

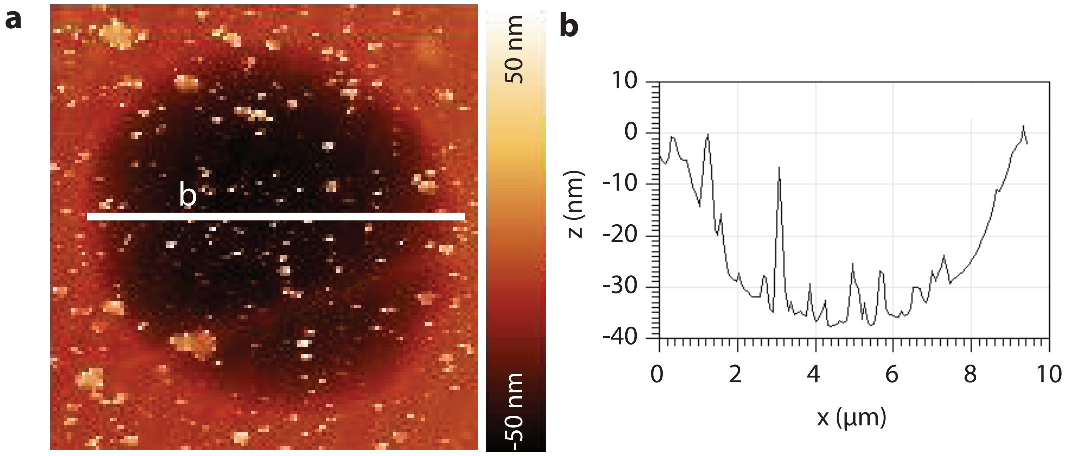


SUPPLEMENTARY FIG. 7. Liquid AFM characterization of a graphene drum a) AFM image of the sample. b) Line scan across the drum, as indicated by a white line in a), shows a maximum deflection of 35 nm.

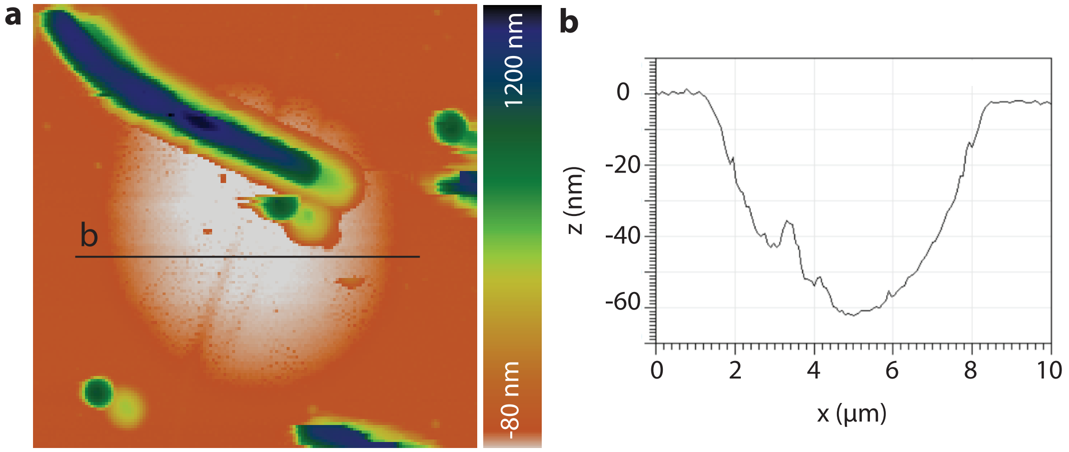


SUPPLEMENTARY FIG. 8. Liquid AFM characterization of a graphene drum with a deposited bacterium a) AFM image of the sample. b) Line scan across the drum, as indicated by a black line in a), shows a maximum deflection of 60 nm.

#### SUPPLEMENTARY NOTE 6: MEASUREMENT ON EXFOLIATED GRAPHENE

For comparison to the CVD bilayer graphene used in all other experiments in this study, we show here a long measurement on an exfoliated graphene flake (thickness <10 nm). The nanomotion was recorded during 10 minutes before administering DNP at a concentration of 2 mM at the point indicated in Figure 9. After a few minutes that are needed for settlement and re-alignment, measurements were continued. After 30 minutes from drug injection, the amplitude of nanomotion begins to decrease. This experiment indicates, that also 2D materials other than CVD bilayer graphene can be suitable candidates for nanomotion detection.

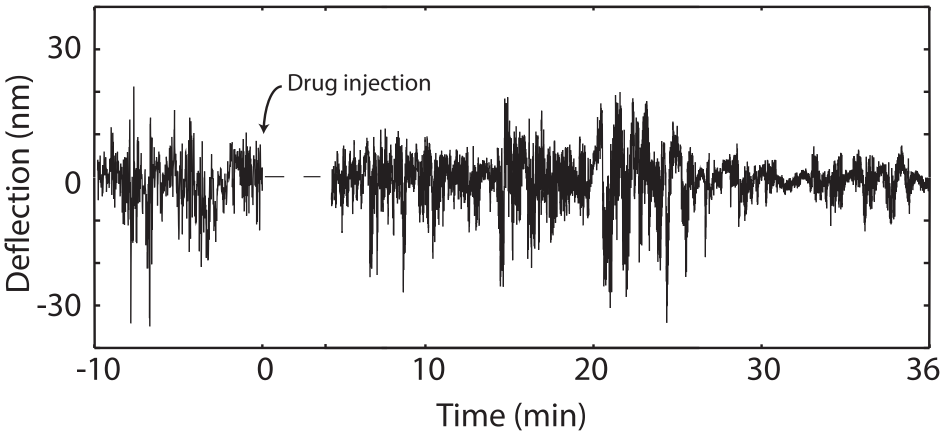


SUPPLEMENTARY FIG. 9. Measurement of *E. coli* nanomotion with exfoliated graphene before and after adding DNP antibiotic.

[1] T. K. Wood, A. F. G. Barrios, M. Herzberg, and J. Lee, Applied microbiology and biotechnology **72**, 361 (2006).
